# A phylogenetic approach to studying the roles of within-host evolution and between-host transmission of resistance for clinical *Escherichia coli* infections

**DOI:** 10.1101/2022.12.16.520823

**Authors:** Florentine van Nouhuijs, MaryGracy Arakkal Antony, Faye Orcales, Landen Gozashti, Scott W. Roy, C. Brandon Ogbunugafor, Pleuni S. Pennings

**Author notes:** These authors contributed equally to this study (Florentine van Nouhuijs, MaryGracy Arakkal Antony, Faye Orcales).

## Abstract

Bacterial antibiotic resistance represents a public health concern that will remain relevant for the foreseeable future. Antibiotic resistant bacterial infections can occur in two ways: (1) a host is infected by a resistant bacterial strain (due to between-host transmission of resistance), or (2) a host is infected infection by a susceptible strain, followed by the de novo evolution or acquisition of resistance (due to within-host evolution of resistance). While both are critical to understanding how the evolution of resistance happens in natural settings, the relative rate at which they occur is unclear. Here, we employ phylogenetic comparative methods to examine the evolutionary dynamics of resistance in *Escherichia coli* for multiple common antibiotics. We report evolutionary patterns consistent with common *de novo* evolution of resistance for some antibiotics and sustained transmission of resistant strains for others. For example, we observe 79 putative *de novo* resistance evolution events for resistance to Cefuroxime but only 31 for resistance to Ciprofloxacin, despite similar numbers of observed infections (239 and 267 respectively). We find that clusters of resistance are generally larger for Ciprofloxacin, Ceftazidima and AmoxiClav, which suggests that for these drugs, resistance is often transmitted from patient to patient. In contrast, we find that cluster sizes for resistance are generally smaller for PipTaz, Cefuroxime and Gentamicin, suggesting that resistance to these drugs is less often transmitted from patient to patient and instead evolves *de novo*. In addition to differences between drugs, we also find that cluster sizes were generally larger in phylogroup B2 compared to the other phylogroups, suggesting that transmission of resistant strains is more common in this phylogroup compared to the others. Our study proposes new approaches for determining the importance of *de novo* evolution or acquisition (within-host evolution) from resistance from infection with an already resistant strain (between-host transmission). Significantly, this work also bridges an important gap between evolutionary genomics and epidemiology, opening up a range of opportunities for studying the evolutionary dynamics of bacterial antibiotic resistance.

## Introduction

Antibiotic resistance is a major global health concern. Each year, two million people in the US are infected by antibiotic-resistant bacteria, and an estimated 23,000 of these die as a result (Centers for Disease Control and Prevention (U.S.) 2019). Further, global estimates put the number of deaths associated with bacterial antimicrobial resistance at 4.95 million in 2019 alone (Murray et al. 2022).

The bacteria that cause resistant infections can arise either because they acquired resistance via new mutation or via horizontal gene transfer, where the genes and mutations responsible for resistance are shared between individual bacterial clones. In both of these cases, the public health problem arises because it is more challenging to treat the individual affected, and because those resistant populations can spread to new hosts, greatly exacerbating the problem. Thus, understanding the evolutionary reasons for the origin and spread of antibiotic-resistant bacterial strains is a major objective in epidemiology (Martínez, Baquero, and Andersson 2007; Davies and Davies 2010; Blanquart 2019; Laxminarayan et al. 2013). Now, let’s consider the same situation from the viewpoint of a single patient (see Figure 1). For a patient, an antibiotic-resistant infection can originate from two sources: one possibility is that the patient originally was infected with a susceptible strain of bacteria, which then evolved to become resistant (within-host evolution or acquired resistance). Alternatively the patient may have been infected by an already resistant strain (between-host spread or transmitted resistance). Note that for a single infection in a single patient, both of these origins may play a role, but for resistance to different drugs. For example, the patient may be infected initially with a strain resistant to Ciprofloxacin (between-host spread or transmitted resistance), and subsequently this same strain evolves to become resistant also to Gentamicin (within-host evolution or acquired resistance).

**Figure 1.**
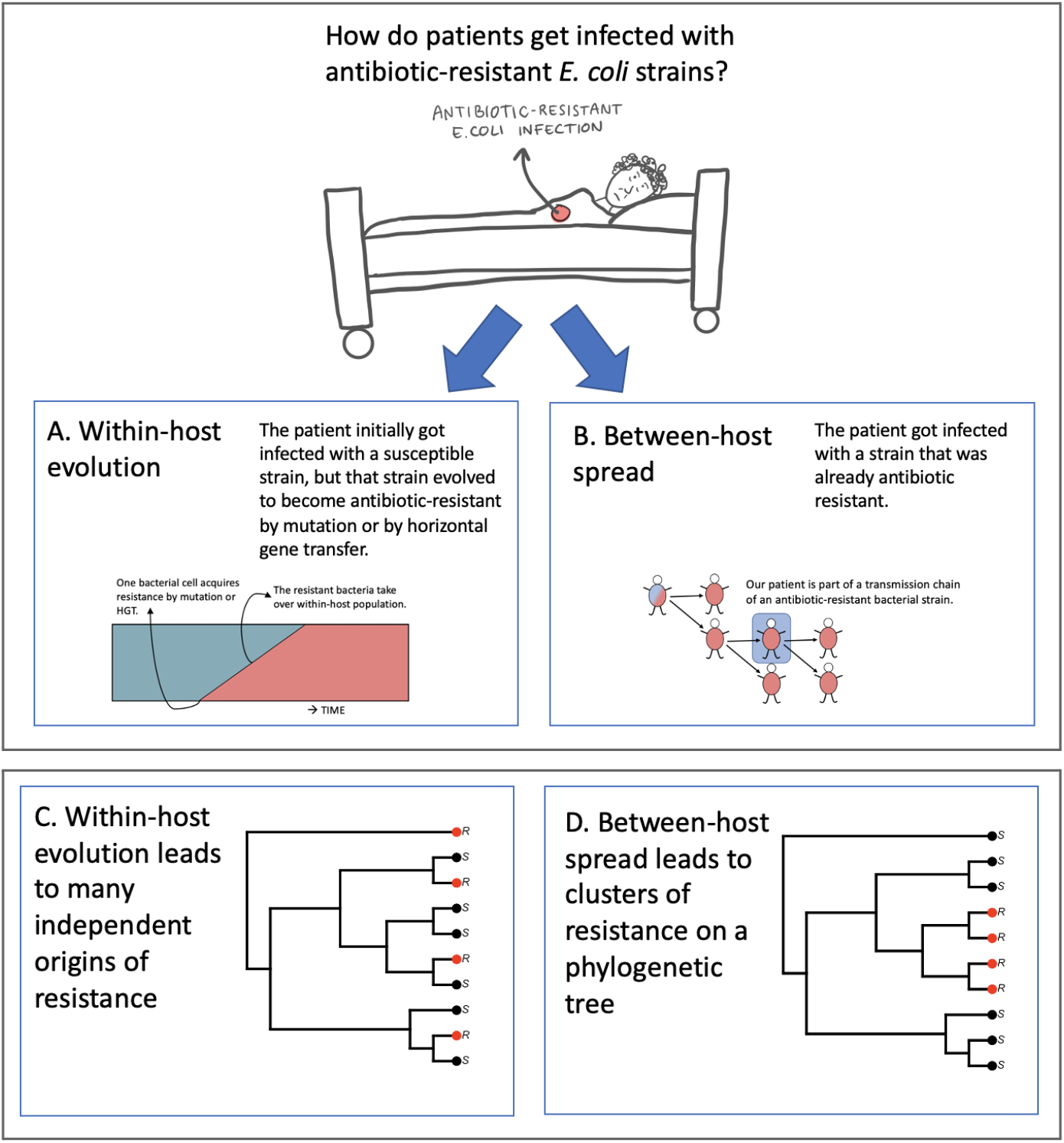
Diagram showing how patients can become infected by antibiotic resistant strains, either through A. within-host evolution or acquired resistance, or through B. between-host spread or transmitted resistance. C. and D. Expectation of what phylogenetic trees will look like depending on whether resistance is typically caused by within-host evolution (C.) or between-host spread of resistance (D.).

For hospitals and healthcare workers, whether the resistance observed at the bedside is (thought to be) arise from *within-host evolution* or *between-host spread* should influence clinical decision-making. For example, if resistance is often transmitted (*between-host spread*), then resistance tests should be carried out before a drug is chosen for treatment. On the other hand, if resistance usually evolves *within the patient*, this could mean that treatments are not optimal, because they allow the evolution of resistance to happen. It would therefore be useful to know, for a given combination of drug and pathogen, how commonly it is due to *within-host evolution or acquired resistance* and how commonly it is due to *between-host spread or transmitted resistance*. Unfortunately, in most situations, we don’t know how common each of these paths are.

The relative role of *within-host evolution* and *between-host spread* of resistance has been studied in several systems, such as HIV (Yang et al. 2015) and *Mycobacterium tuberculosis* (Kendall, Fofana, and Dowdy 2015; Knight et al. 2019). For these two pathogens it is known that both *within-host evolution* and *between-host spread* are important drivers of resistance, although the relative roles of each are debated (for *M. tuberculosis*) and have changed over time (with *within-host evolution* losing importance for HIV over the years (Yang et al. 2015)). When we consider other pathogens, there are examples where one or the other mechanism clearly dominates. For example, malarone resistance in the malaria parasite *Plasmodium falciparum* can evolve within hosts but appears to be unable to transmit to other hosts. We know this because malarone resistance has been observed in patients treated with malarone, but never in patients not on the drug (Musset, Le Bras, and Clain 2007). Apparently (and lucky for us), a *Plasmodium falciparum* strain with malarone resistance cannot be transmitted from person to person, which makes that malarone resistance is still very rare even after many years of use, and malarone remains a recommended drug for all areas where *P. falciparum* is found according to CDC. The opposite situation also is found in *P. falciparum*: there are drug-resistant strains that have spread world-wide because they are so easily transmitted from person to person (Mita, Tanabe, and Kita 2009; Roper et al. 2004). Another example of extensive between-host spread of resistant strains is found in MRSA (methicillin resistant *Staphylococcus aureus*), where the USA300 strain has spread across all continents (Strauß et al. 2017; Laxminarayan et al. 2013). For many other pathogens, we expect that both within-host evolution (including by HGT) and between-host spread of resistant strains are important factors that contribute to the burden of resistant infections.

Our overarching goal is to determine the importance of within-host evolution and between-host spread for drug resistance in pathogens. While different approaches to tackle this big question have been used (Yang et al. 2015; Pradhananga et al. 2022), here we will use a phylogenetic approach. There is a long tradition of using phylogenetic trees to study the evolution of pathogens with so-called phylodynamic methods (Volz, Koelle, and Bedford 2013). For example, phylodynamic studies have been used to study immune escape in influenza and to understand transmission patterns in HIV (Volz, Koelle, and Bedford 2013). Phylodynamic approaches have also been used to study aspects of drug resistance (Kühnert et al. 2018; Pečerska et al. 2021).

We propose that patterns of phylogenetic clustering can be used to determine the relative importance of within-host evolution and between-host transmission for resistance to different drugs. Clustering patterns contain this kind of information because if resistance to a drug evolves within a host and is typically not transmitted to other hosts, the tree should be characterized by resistance on separate individual leaves (Figure 1C). On the other hand, if resistance to a drug can easily be transmitted, we should see evidence of that on the phylogenetic tree as reflected by clustering of resistant strains (Figure 1D).

It would be of interest to determine whether clustering patterns differ between different pathogens, but instead, we decided to first investigate a simpler question: for the same pathogen does clustering differ between resistances to different drugs? In other words, given one tree, does resistance to drug A occur in a more clustered fashion than resistance to drug B? If this were the case, then we’d know it is not because of sampling issues, because we are looking at exactly the same samples, and instead it would likely be due to differences in the probability of transmission of the resistant strains.

To be able to compare resistances to different drugs, we searched for a large genomic dataset for which resistance phenotypes were available for several drugs. We found such a dataset in a study on *E coli* bloodstream infections (bacteremia) in the UK (Kallonen et al. 2017). *E. coli* is a common cause of bloodstream infections (de Kraker et al. 2013; Peralta et al. 2007; Elixhauser, Friedman, and Stranges 2006; Gerver et al. 2015) and the incidence of multidrug-resistant (MDR) *E. coli* and extended-spectrum β-lactamase (ESBL)-producing *E. coli* that cause bloodstream infections are on the rise (Gladstone et al. 2021).

The Kallonen dataset consists of *E. coli* genome sequences from 1506 bacteremia patients with phenotypic information on resistance to 6 drugs: Cefuroxime, AmoxiClav, Ciprofloxacin, Gentamicin, PipTaz and Ceftazidime. In the original dataset there is information about other drugs too, but there is a lot of missing data for these drugs and we therefore decided to not include them in our study. E. coli is typically split in several phylogroups. We consider here A, B1, B2 and the combination of D and F.

For each drug, and for each *E. coli* phylogroup we determine the cluster sizes of drug resistance on the phylogenetic tree and we then compare whether resistance to any of the drugs is associated with a different distribution of cluster sizes. We find that cluster sizes are typically larger for Ciprofloxacin, Ceftazidime and AmoxiClav, whereas cluster sizes are typically smaller for PipTaz Cefuroxime and Gentamicin. This provides evidence that between-host transmission is relatively more important for Ciprofloxacin, Ceftazidime and AmoxiClav when compared to PipTaz Cefuroxime and Gentamicin.

Our results show that cluster sizes do indeed differ between resistances to different drugs, which is consistent with different roles of within-host evolution and between-host transmission between the different resistances. This, in turn, suggests that prevention of resistance may require different strategies for these different drugs. We have hope that this study and the approach will lead to a better understanding of the causes of drug resistance and potential ways to prevent antibiotic resistant infections.

## Methods

### Accessing relevant data

The data that is used is obtained from a study on *E. coli* in the UK sampled from patients with bacteremia (Kallonen et al. 2017). The data contains WGS data for 1509 *E. coli* isolates obtained from two different collections. The first collection contains 1094 isolates that are collected between 2001-2011 by 11 hospitals across England. The second collection contains 415 isolates from a diagnostic laboratory at the Cambridge University Hospitals NHS Foundation Trust in Cambridge stored between 2006 and 2011. All the isolates are associated with bacteremia.

We downloaded sequence reads for all samples considered in this study from the NCBI Sequence Read Archive (SRA) using fasterq-dump (Leinonen, Sugawara, and Shumway 2011). See Supplementary Table 1 (S1_Phylogroup.csv) for sample accession numbers. We also retrieved corresponding metadata for all considered samples directly from (Kallonen et al. 2017).

All data and code is available on https://github.com/FlorentinevanNouhuijs/Team_phylo.

### Core genome alignment, SNP calling and filtering

We used the *Snippy* pipeline (https://github.com/tseemann/snippy) with default parameters to generate a core genome alignment. *Snippy* employs *Freebayes* (Garrison and Marth 2012) to call SNPs for each sample against the core genome reference, then constructs multiple sequence alignments of core genome sequences across all samples. We filtered poor quality genotypes and sample-specific zero coverage positions from our core alignment using *snippy-clean_full_aln* with default parameters. Then, we employed *Gubbins* (Croucher et al. 2015) with default parameters to remove recombinant sequences from our alignment and produce a final cleaned alignment file for SNP calling. We employed *snp-sites* (Page et al. 2016) with default parameters to call SNPs from our final cleaned core genome alignment.

### Phylogenetic reconstruction

We used *IQ-TREE* (Minh et al. 2020) with a general time reversible model with unequal rates and unequal base frequencies (GTR) model to reconstruct a phylogeny across all samples with 1000 bootstraps. Considered *E. coli* samples generally represent five phylogenetically distinct clades, or phylogroups. Internal nodes demonstrate strong bootstrap support suggesting confident reconstruction of *E. coli* phylogroups. Concomitant to this result, we observe clear clustering of phylogroups on our tree with the exception of phylogroups D and F, which have been previously shown to be paraphyletic (Kallonen et al. 2017). For the analyses in this study, we therefore combined phylogroups D and F into one tree.

### Extracting subtrees for each phylogroup

After generating a global phylogeny across all samples, we extracted subtrees for each phylogroup using the *phytools* package in R (Revell 2012). To ensure that the trees were bifurcating, we resolved any polytomies by using the ape package’s *multi2di* function (Paradis and Schliep 2019).

### Determining cluster sizes

To explore evolutionary patterns of *E. coli* antibiotic resistance we investigated cluster sizes of resistant samples across different antibiotics for each phylogroup. To do this we first simulated a single history of the resistance phenotype on the tree using the *make*.*simmap* function from *phytools* (Revell 2012). Next, for each resistant sample, we traversed up the tree until we found a node which was not resistant. Once we reached such a node, we determined the resistance cluster size based on the number of daughters for the previous node. If a sample did not have a resistant neighbor, we assigned it a cluster size 1 (see Figure 2 for an example).

**Figure 2.**
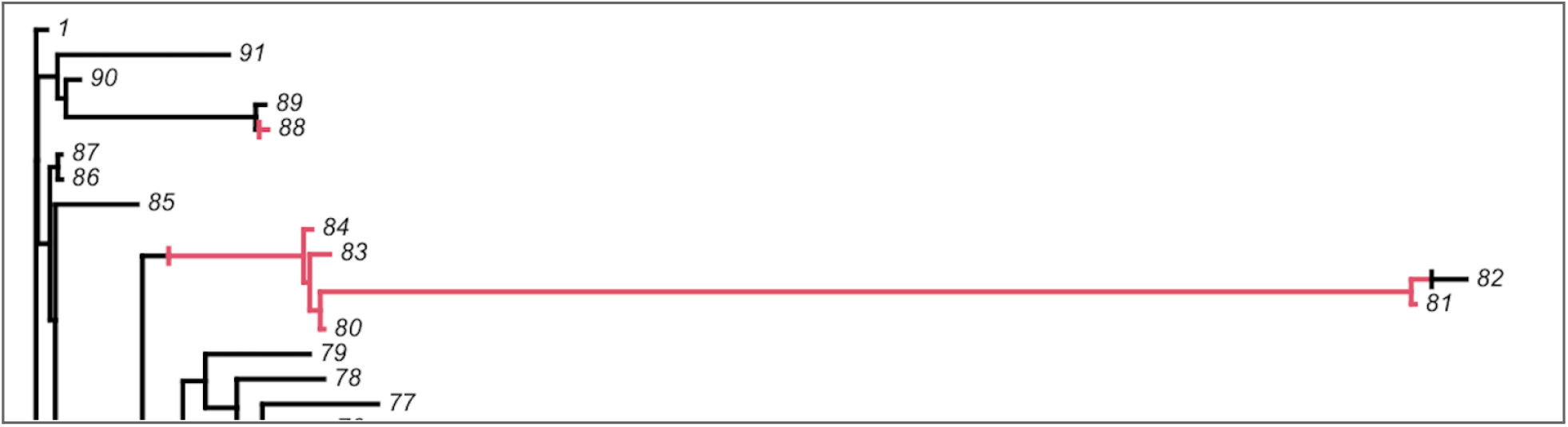
Example cluster size calculation. Here, tip number 88 is a cluster of size 1. Tip 84, 83, 81 and 80 make up a cluster of size 4. Red: resistant, black: susceptible.

### Statistical analysis

First, for each drug and each phylogroup (A, B1, B2, D&F), we visualized the subtree to inspect the distribution of resistant strains on the trees. For all analysis, we grouped phylogroups D and F because they are paraphyletic (Kallonen et al. 2017). Next, after determining a list of cluster sizes for each of 6 drugs and for each phylogroup, we plotted the distribution of cluster sizes for each drug. We then used a Mann-Whitney U test for each pair of drugs to determine whether the cluster size distribution differed between the drugs. Next, we plotted the fraction resistant against the average cluster size for each phylogroup separately and used a linear model to determine what best explains the fraction of resistant samples, using drug and average cluster size as explanatory variables. Next, we fitted a generalized linear model (with quasi-poisson error family) to see which factors best explain the observed cluster sizes. We find that a model with both drug and phylogroup as factors (but not their interaction) best explains the observed cluster sizes.

We also used a permutation-based approach to investigate whether resistance phenotypes clustered on the tree more than expected by chance. To do this, we used *phyloclust* from the R package *RRphylo* (Castiglione et al. 2018) with the parameter *nsim=1000. phyloclust* computes the mean cophenetic distance between all samples under the focal state (resistance in our case), and compares this distance to a random distribution of distances obtained by sampling as many random tips as those under the focal state *nsim* times.

## Results

An overview of the samples we used for our analysis is given in table 1.

**Table 1.**
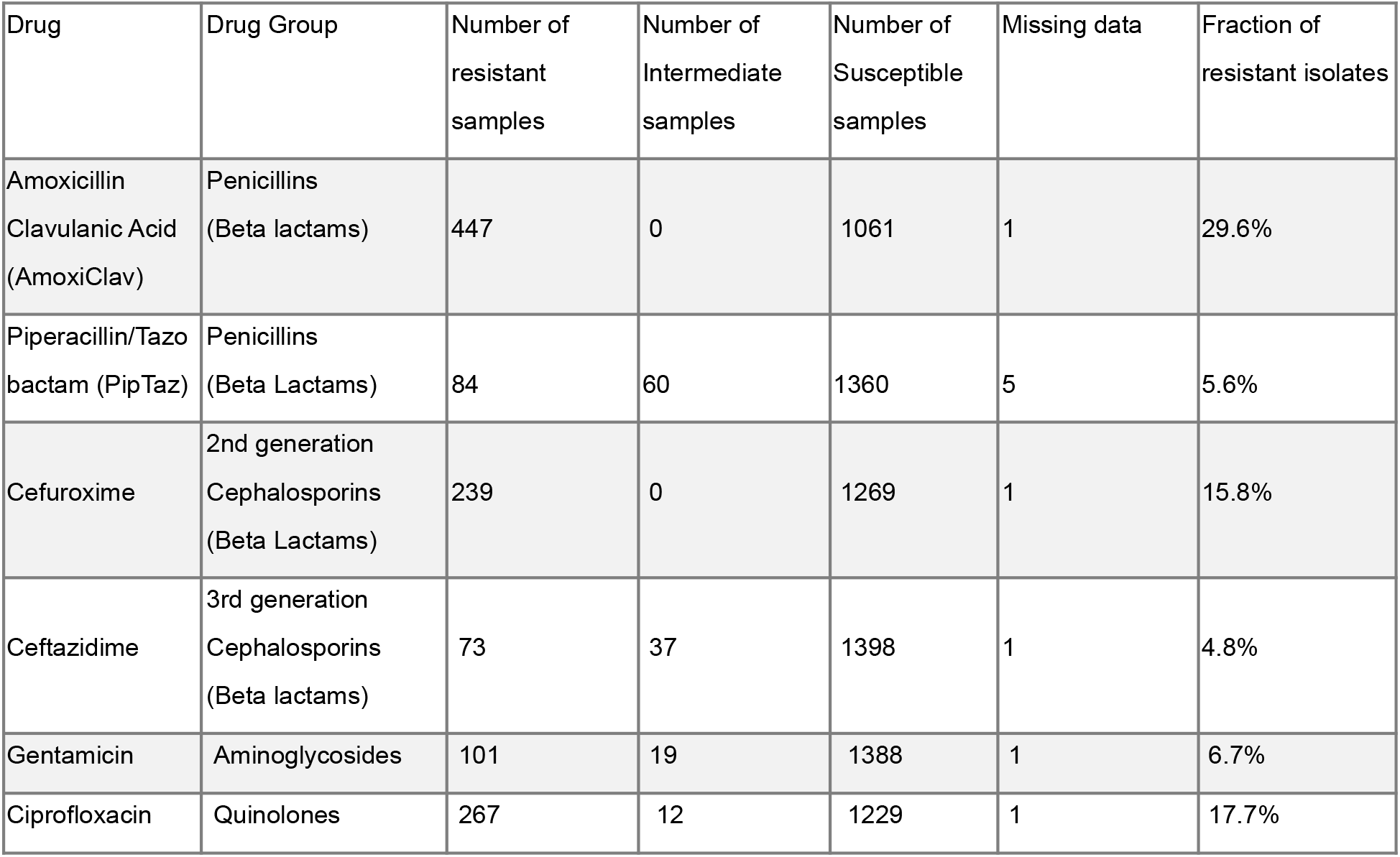
Overview of samples in our dataset

| Drug | Drug Group | Number of resistant samples | Number of Intermediate samples | Number of Susceptible samples | Missing data | Fraction of resistant isolates |
| --- | --- | --- | --- | --- | --- | --- |
| Amoxicillin<br>Clavulanic Acid<br>(AmoxiClav) | Penicillins<br>(Beta lactams) | 447 | 0 | 1061 | 1 | 29.6% |
| Piperacillin/Tazo<br>bactam (PipTaz) | Penicillins<br>(Beta Lactams) | 84 | 60 | 1360 | 5 | 5.6% |
| Cefuroxime | 2nd generation<br>Cephalosporins<br>(Beta Lactams) | 239 | 0 | 1269 | 1 | 15.8% |
| Ceftazidime | 3rd generation<br>Cephalosporins<br>(Beta lactams) | 73 | 37 | 1398 | 1 | 4.8% |
| Gentamicin | Aminoglycosides | 101 | 19 | 1388 | 1 | 6.7% |
| Ciprofloxacin | Quinolones | 267 | 12 | 1229 | 1 | 17.7% |

### Visualizing subtrees

Antibiotic resistant samples show diverse evolutionary patterns across drugs and phylogroups. First, for each drug and each phylogroup, we visualized the subtree to inspect the distribution of resistant strains on the trees. Phylogroup D and F were merged because they are paraphyletic (Kallonen et al. 2017). In figure 3A, we show that in Phylogroups D and F, Ciprofloxacin resistance occurs mostly in clusters. In this subtree, there are 42 Ciprofloxacin resistant samples that cluster in 9 clusters. There are 4 clusters of size 1 and 5 clusters with more than one sample (size 2, 3, 7, 8, and 18). In figure 3B we show that in the same phylogroups (D and F), resistance to another drug (Gentamicin) is much less clustered. In this case, there are 14 resistant samples, in 13 clusters (12 clusters of size 1, and one cluster of size 2). This suggests that in phylogroups D and F, both Ciprofloxacin and Gentamicin resistance have evolved several times (at least 9 and 13 times), but Ciprofloxacin resistance can spread from patient to patient more easily than Gentamicin resistance, leading to more Ciprofloxacin-resistant infections (42) than Gentamicin-resistant infections (14).

**Figure 3.**
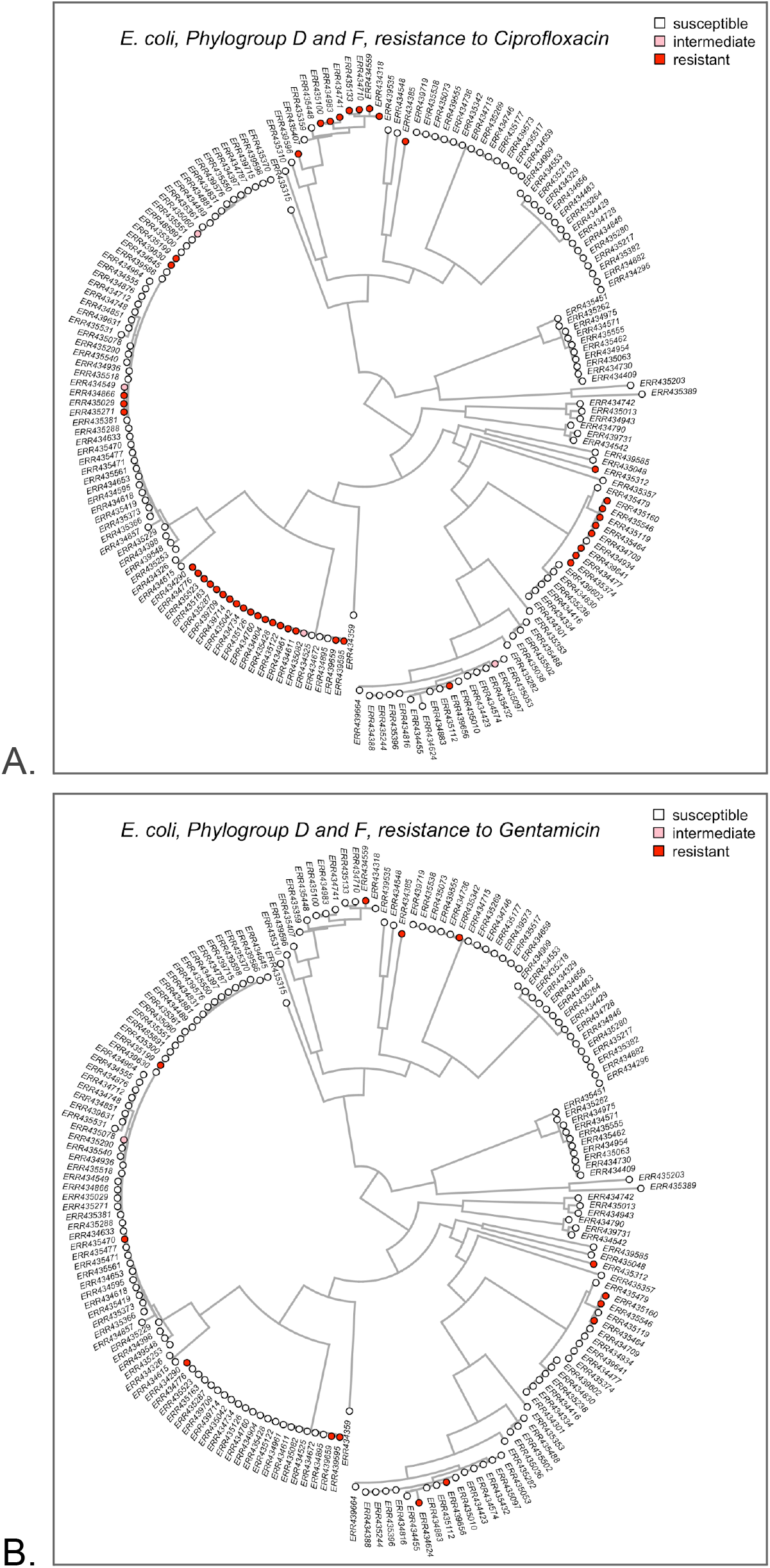
A. Ciprofloxacin resistance in Phylogroups D and F occurs mostly in clusters. B. Gentamicin resistance in Phylogroups D and F occurs mostly on single tips of the tree.

### Cluster size distributions

When drug resistance is transmitted often from one patient to another, it leads to clusters of resistance on the phylogenetic tree. This means that cluster sizes can inform us about the relative importance of transmission of resistance. We started by making a list containing cluster sizes for each cluster of each of the 6 drugs (Cefuroxime, AmoxiClav, Ciprofloxacin, Gentamicin, PipTaz and Ceftazidime). Figure 4A shows the total number of clusters for each of the drugs. The total number of clusters is of interest because it provides an estimate of the number of origins of resistance in our dataset (Kreiner et al. 2019). We notice here that there are many clusters for each of the drugs, ranging from 24 for Ceftazidime to 134 clusters for AmoxiClav resistance.

**Figure 4A.**
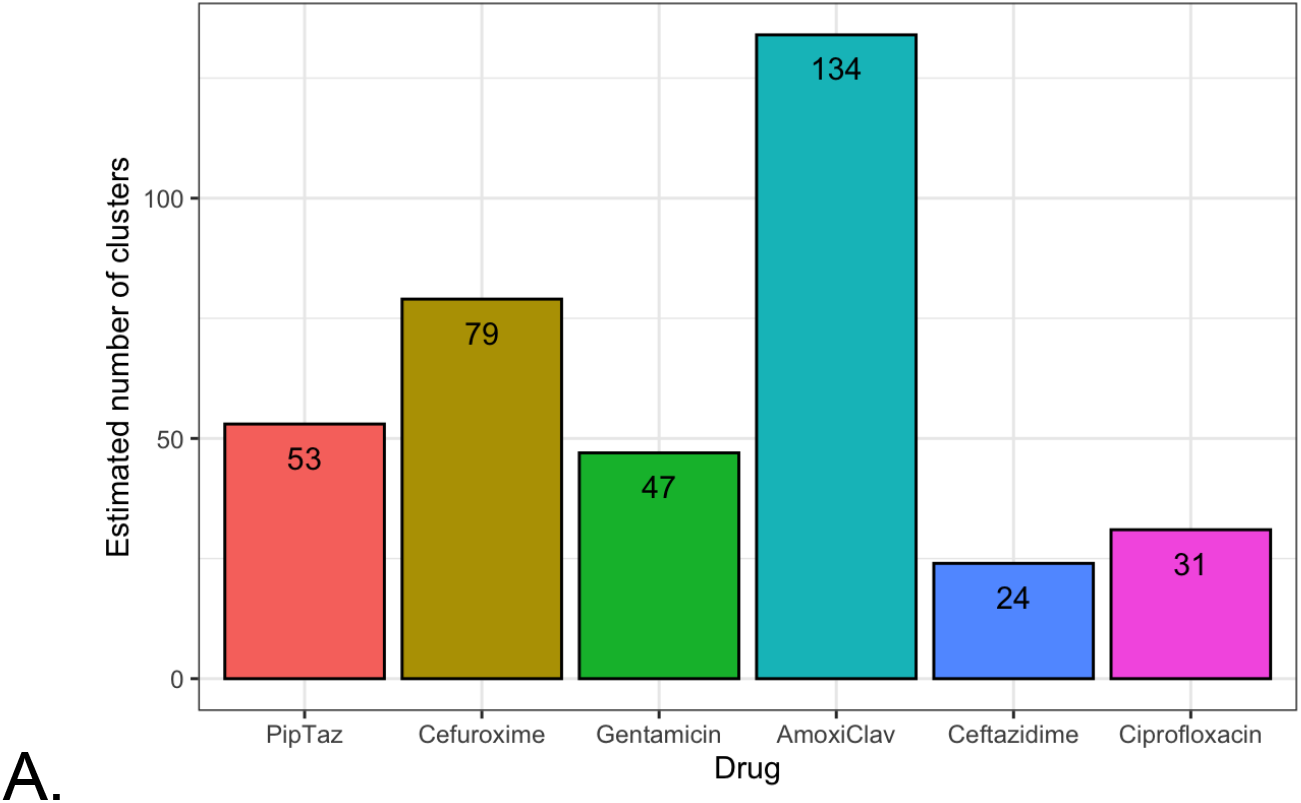
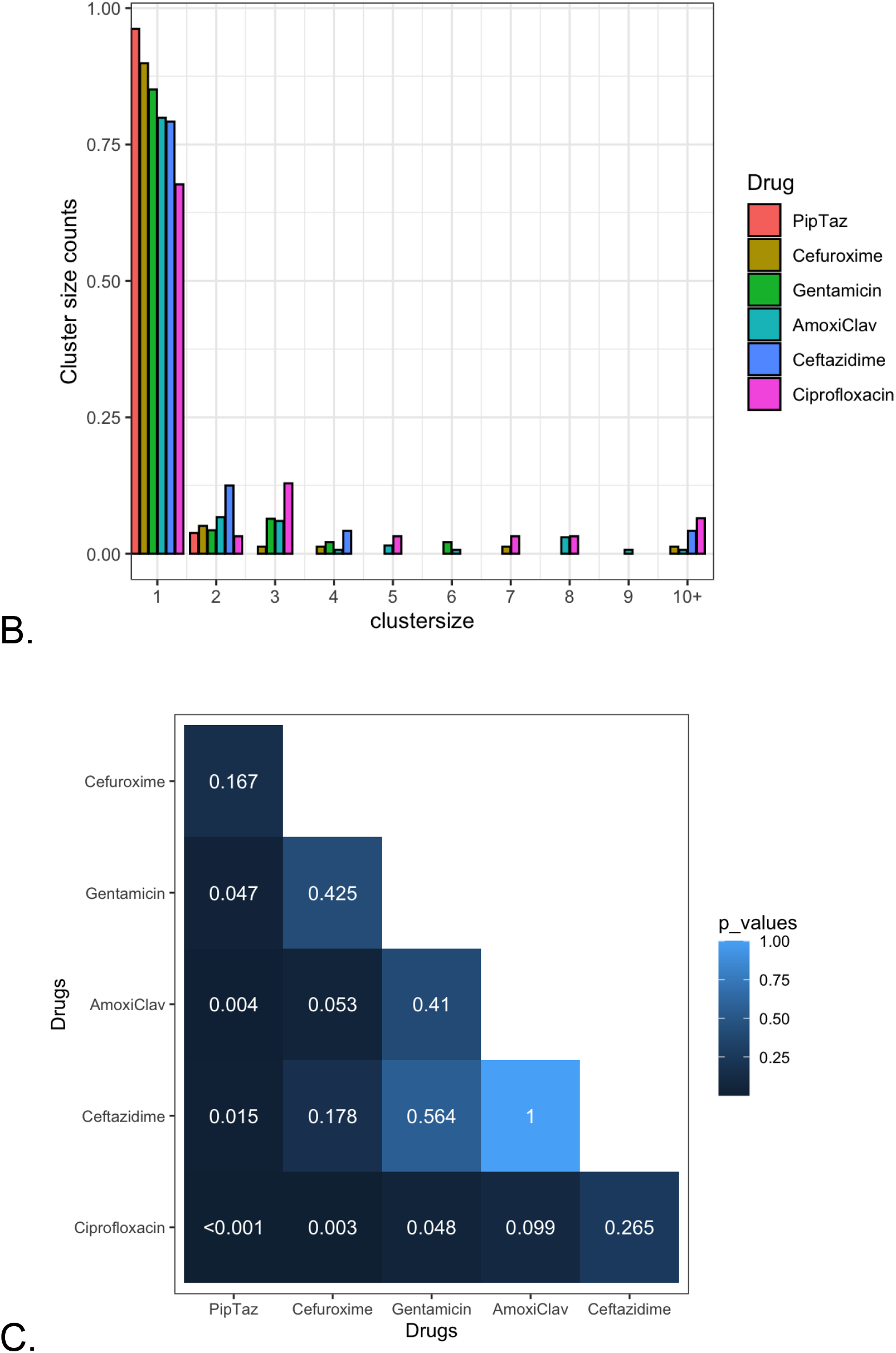
Total number of clusters for resistance to Cefuroxime, AmoxiClav, Ciprofloxacin, Gentamicin, PipTaz and Ceftazidime. B. Distribution of cluster sizes for resistance to Cefuroxime, AmoxiClav, Ciprofloxacin, Gentamicin, PipTaz and Ceftazidime. C: Heatmap of p-values from Mann-Whitney U tests for each pair of drugs to determine whether the cluster size distribution differed between the two drugs.

Figure 4B shows the distribution of cluster sizes for each of the 6 drugs. We find that most clusters (309 of 368 clusters) were size one. The other 59 clusters that are larger than size one contain more than half of resistant samples (438 out of 747 samples cluster in 59 clusters). In Figure 4B, we can see that there are differences in the cluster size distributions, with, for example, larger cluster sizes for Ciprofloxacin and smaller cluster sizes for PipTaz. We used a Mann-Whitney U test for each pair of drugs to determine whether the cluster size distribution differed between the two drugs (Figure 4C). We find that 6 out of 30 comparisons were significant at the 0.05 level. Specifically, the distribution of cluster sizes for Ciprofloxacin resistance was significantly different than the distribution of cluster sizes for Cefuroxime, Gentamicin, and PipTaz. The distribution of cluster sizes for PipTaz was different from that of Ciprofloxacin, Ceftazidime, AmoxiClav and Gentamicin.

### Relationships between cluster size, phylogroup, drug and fraction resistant samples

When clusters are large, they affect many samples. We therefore hypothesize that the fraction of resistant samples in a phylogroup could be explained by the average cluster size. However, when we fit a linear model to the fraction of resistant samples, using drug and average cluster size as explanatory variables, we see that several drugs have a significant effect, but the average cluster size doesn’t (Figure 5). Specifically, when PipTaz is used as the reference, there is a significant positive effect (more resistance) for Cefuroxime, AmoxiClav and Ciprofloxacin.

**Figure 5:**
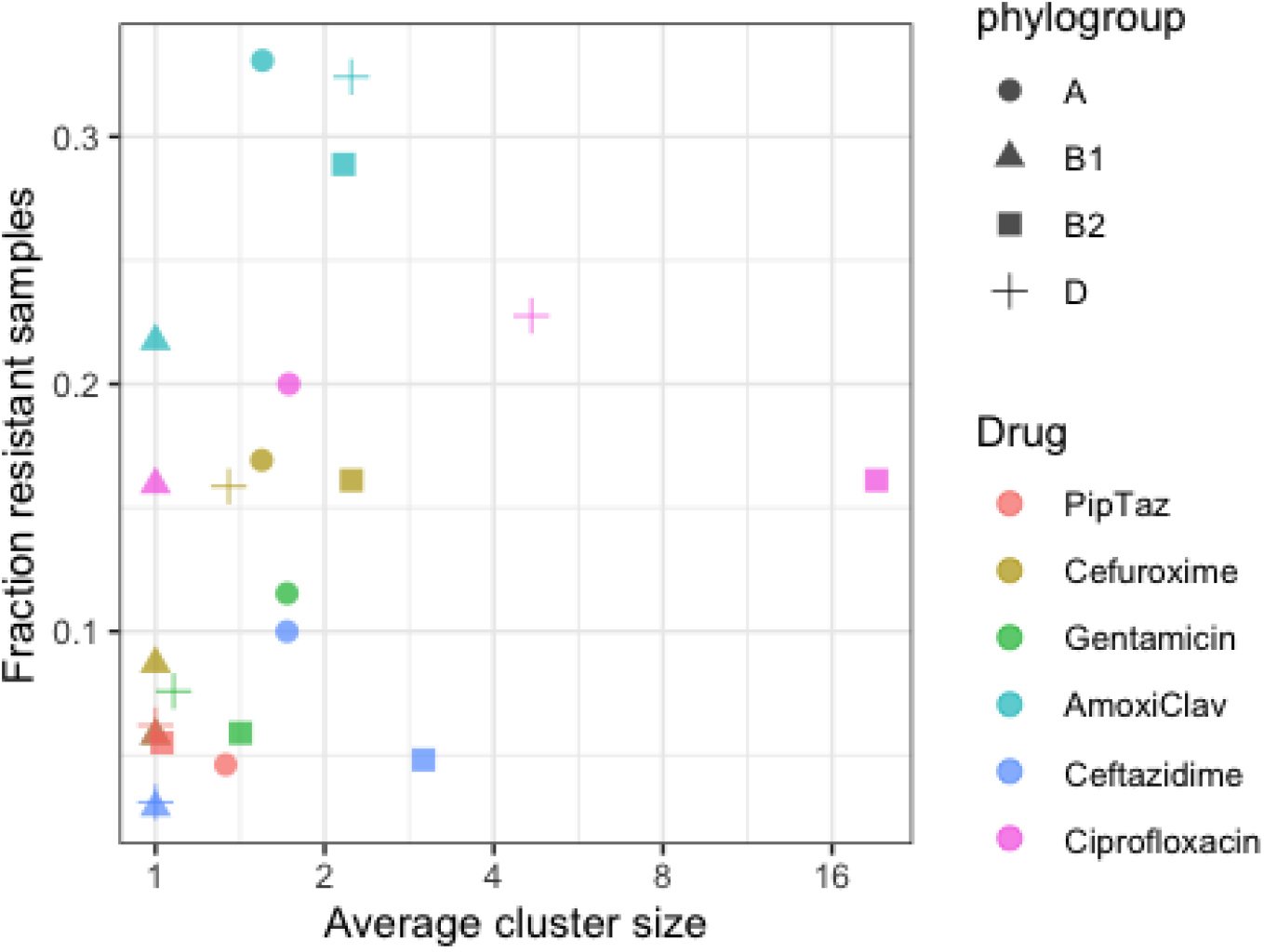
The average cluster size versus the fraction of resistant samples for each phylogroup and each drug.

Next, we fitted a generalized linear model (with quasi-poisson errors) to see which factors best explain the observed cluster sizes. We find that a model with both drug and phylogroup as factors (but not their interaction) best explains the observed cluster sizes, with significantly larger cluster sizes in phylogroup B2 and for Ciprofloxacin.

### Phyloclust analysis

While comparison of resistant cluster size distributions represents a simple method for differentiating between modes of resistance evolution across different antibiotics, this approach only relies on topological relationships among samples and does not consider their phylogenetic relatedness. In light of this, we also used a complementary approach to test for phylogenetic clustering (or lack thereof) of resistant samples, which considers distances between samples. For each phylogroup and drug, we compared the observed average cophenetic distance (that is, the sum of branch lengths between two samples) between resistant samples to expectations from randomization (see Methods). We find that our results differ in some ways from the cluster size approach. We find significant p-values for the *phyloclust* test only in phylogroup B2, and don’t find significant results in cases where we had observed fairly large cluster sizes (e.g., for Ciprofloxacin resistance in group D/F, see figure 3A and 4), see Figure 6A. These results mean that resistant samples in phylogroup B2, for most drugs, are more closely related to each other than expected if they were randomly distributed on the tree, but this is not the case in the other phylogroups. This is the case because in phylogroup B2 there is only one main cluster with most of the resistance (Figure 6B), whereas in the other phylogroups, at least for some of the drugs, we see multiple significant, but smaller clusters of resistance (Figure 3A).

**Fig 6A.**
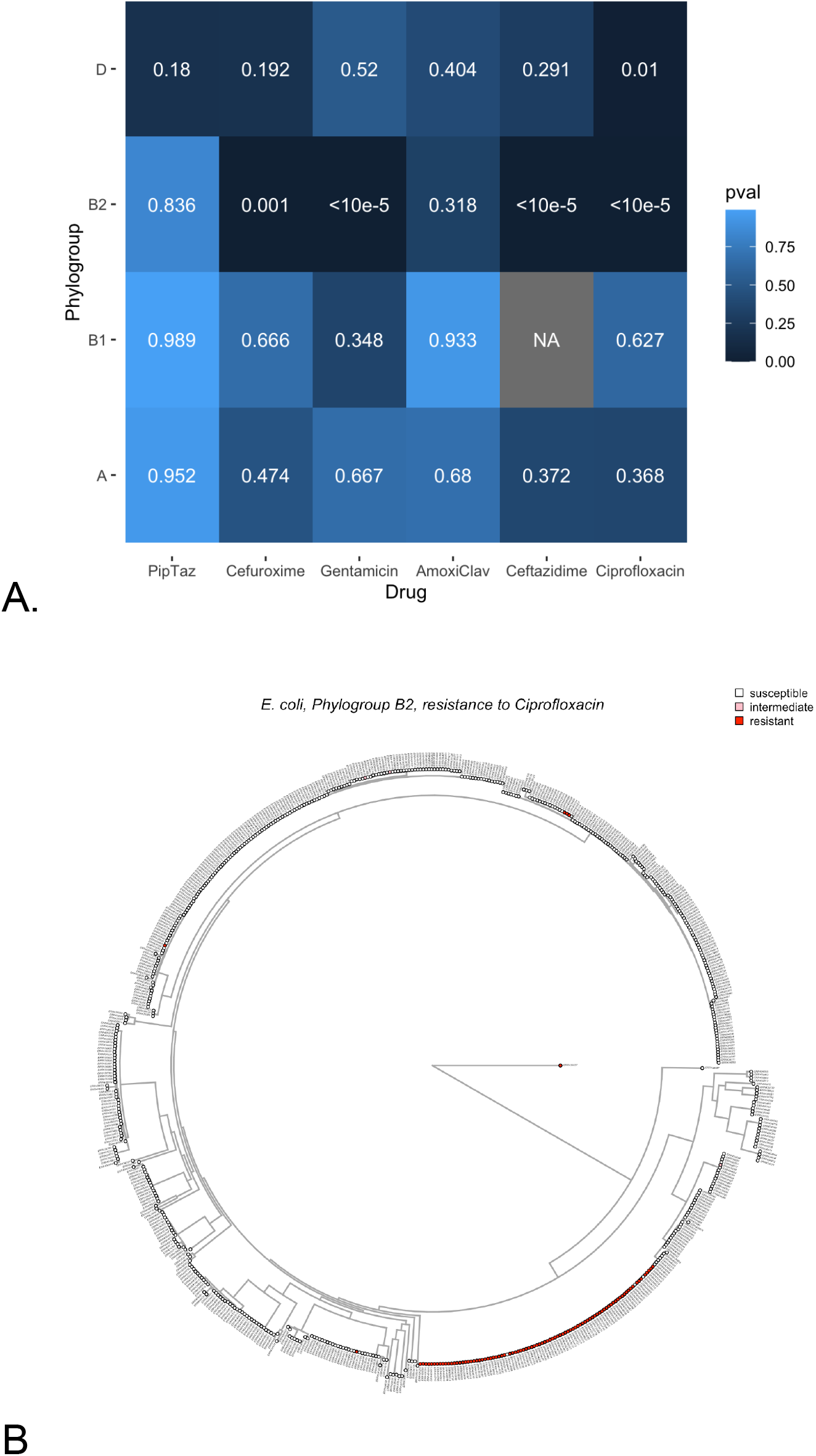
Phyloclust test p-values for all drugs and phylogroups. B. Phylogenetic tree for Ciproflxacin resistance in phylogroup B2, where 90 resistant samples are found in one cluster. 6 resistant samples are found in 4 other clusters.

## Discussion

In this study, we aimed to use evolutionary theory and comparative phylogenetic methods to understand the dynamics of the evolution of resistance in clinical samples of resistant bacteria. To do this, we analyzed an existing large genomic *E. coli* dataset from the United Kingdom containing over 1500 *E*.*coli* samples, taken over the course of a decade, from hospitals across England (Kallonen et al. 2017). We focused on this particular dataset because of its size (over 1500 genomes) and the fact that it had near complete phenotypic data for resistance to 6 different drugs (Table 1). This combination of genomes and phenotypes allowed us to study drug resistance clustering in a phylogenetic context. We analyzed each of four subtrees separately: phylogroups A, B1, B2 and D/F. D and F are paraphyletic and for our analysis it made more sense to study them together as one phylogroup. For each of the four subtrees, we simulated the history of the resistance phenotypes on the tree. We then counted, for each origin of resistance, how many tree tips were affected, which we call the cluster size, and we analyzed the distribution of cluster sizes.

We are interested in the number of tree tips that are affected by a single origin of resistance because it can be seen as a measure of how often resistant strains are transmitted from one patient to another. If a single origin of resistance leads to many resistant tree tips, it means that between-host transmission of the resistant strain must have occurred many times. On the other hand, if a single origin of resistance only leads to a single resistant tree tip, then between-host transmission may not have occurred at all.

It should be noted here that with the methods we used, we cannot quantify how common or uncommon between-host transmission of resistance is. This is because we don’t know how many patients were involved in a transmission chain, yet not sampled. For example, if our dataset consists of 1% of all *E. coli* infected patients, then a cluster of size 10 may, in reality, be of size 1000, and a cluster of size one may, in reality, be of size 100. However, if there is no between-host transmission of resistance, then no matter how many patients are sampled, the cluster size would still be one – with increased sampling, we would just recover more clusters of size one. We hope that at some point there will be mathematical models available that allow us to infer rates of between-host transmission of resistant strains from observed cluster size distributions. For example, it may be feasible to add a phylogenetic layer to existing SIR type models of drug resistance evolution (Lipsitch, Bergstrom, and Levin 2000). For now, we focus on the relative differences in cluster sizes between drugs, given that the sampling intensity is the same.

For each of the 6 drugs, we found many independent clusters, the smallest number was 24 (Ceftazidime) and the largest 134 (AmoxiClav). This shows that resistance evolution has occurred many times for each of the drugs, but there are big differences as well, including substantial differences in the number of origins of resistance for drugs that have similar fractions of resistant samples. For example, we observe 79 putative *de novo* resistance evolution events for resistance to Cefuroxime but only 31 for resistance to Ciprofloxacin, despite similar numbers of resistant samples in the dataset (239 and 267 respectively). This difference is consistent with a model where between-host transmission is much more common for Ciprofloxacin than for Cefuroxime.

Our main result is that cluster sizes are generally larger for Ciprofloxacin, Ceftazidime and AmoxiClav and generally smaller for PipTaz, Cefuroxime and Gentamicin. This suggests that between-patient transmission is relatively more important for resistance to Ciprofloxacin, Ceftazidima and AmoxiClav when compared to PipTaz, Cefuroxime and Gentamicin. We also found that clusters were larger in phylogroup B2 compared to the other phylogroups. This may be driven by ST 131, a sequence type that is known for high levels of resistance that came up in the 2000s (Kallonen et al. 2017; Murray et al. 2022; Blanquart 2019).

Finally, we used a separate approach to test for clustering of resistance phenotypes on the tree using the *phyloclust* function from the *RRphylo* package (Castiglione et al. 2018). Here we tested whether the resistance phenotypes clustered on the tree more than expected by chance. We found that this was the case only for some of the drugs and mostly in phylogroup B2. This is the case because almost all resistance in B2 is due to a single large cluster, whereas resistance in the other phylogroups is due to several clusters. The situation in phylogroup B2 could be described as a single-origin incomplete hard sweep, whereas the situation in the other phylogroups could be described as multiple-origin soft sweeps (Hermisson and Pennings 2017). Figure 6B shows the clustering of Ciprofloxacin resistance in phylogroup B2 where almost all resistance is found in just one clade. This clade is the well-known sequence type ST131 which arose in the 2000s (Kallonen et al. 2017). In future studies it would be of interest to determine why there is only one main resistance cluster in the large B2 phylogroup, compared to the other phylogroups.

Antibiotic-resistant infections can be caused by within-host evolution and between-host spread of resistant strains. There is currently no standard method to quantify the role of these two processes. However, with our comparative phylogenetic analyses we were able to show that the roles are not equally important for all drugs. Between-host spread is more important (relatively speaking) for Ciprofloxacin, Ceftazidime and AmoxiClav. Within-host evolution is more important for PipTaz, Cefuroxime and Gentamicin. PipTaz is the most extreme case, as there are 53 origins of resistance in our dataset, and 51 of these origins only affect a single sample (a cluster size of 1), whereas only two origins affect two samples (only 4% of clusters are larger than size 1). This suggests that strains resistant to PipTaz are unlikely to be transmitted to other patients, possibly because of fitness costs of the genes or mutations that cause resistance (Pennings, Ogbunugafor, and Hershberg 2020; Melnyk, Wong, and Kassen 2015; Andersson and Hughes 2011; Helekal et al. 2022). On the other extreme, Ciprofloxacin resistance is often found in sizable clusters. Out of 31 origins of Ciprofloxacin resistance, 21 affect just one sample, while 10 affect more than 1 sample (32% of clusters are larger than size 1). The result is that just 31 origins of resistance to Ciprofloxacin affect 267 resistant samples. Thus, while PipTaz resistance has evolved more often in this dataset, Ciprofloxacin resistance affects many more samples because of between-host transmission.

One possible way to think about our results is to consider the public health risk of using a particular drug for a patient. Our results suggest that *E. coli* strains with resistance to Ciprofloxacin, Ceftazidime or AmoxiClav can spread to other patients, whereas *E. coli* strains with resistance to PipTaz, Cefuroxime or Gentamicin do not often spread to other patients. This could mean that it is more important to prevent evolution of resistance to Ciprofloxacin, Ceftazidime or AmoxiClav, as it could potentially affect many other patients in addition to the one being treated. Ideally, for each combination of drug and pathogen, we could not only know the risks and benefits of using the drug as treatment, but also the potential risk of starting a transmission chain spreading drug resistance to other patients.

A major limitation of our approach is that we have studied just one dataset from one country, focusing on patients with one diagnosis (bacteremia). While this dataset is large (more than 1500 patient samples), it is also a convenience sample from different locations and different years. Future studies will show whether these results hold when we study other *E. coli* datasets and other pathogens.

Our overall goal is to quantify the roles of within-host evolution and between-host spread for drug-resistant infections. We found clear differences in phylogenetic clustering for resistance to different drugs in a large *E. coli* dataset. We hope that this work will inspire studies on other datasets and mathematical modeling approaches that will make it possible to determine the rates of within-host evolution and between-host spread for any combination of drug and pathogen. This will ultimately help us determine whether, for a given drug and pathogen, prevention strategies should focus on within-host evolution or on between-host spread.

## Acknowledgements

We would like to thank Drs Julia Kreiner and Ruth Hershberg for many discussions. This work was supported by NSF grant 1655212 including a generous year-long research experience supplement for Faye Orcales. CBO acknowledges support from National Science Foundation’s Department of Environmental Biology Award Number 2142719 and from the MLK Jr Visiting Scholars Program at the Massachusetts Institute of Technology.

